# Are Granulins Copper Sequestering Proteins?

**DOI:** 10.1101/2020.07.24.220665

**Authors:** Anukool A. Bhopatkar, Vijayaraghavan Rangachari

## Abstract

Granulins (GRN 1-7) are short (∼6 kDa), cysteine-rich proteins that are generated upon the proteolytic processing of progranulin (PGRN). These modules, along with their precursor, have been implicated in multiple pathophysiological roles, especially in neurodegenerative diseases. Our previous investigations into GRN-3 and GRN-5 reveal them to be fully disordered in the reduced form and implicate redox sensitive attributes to the proteins. Such redox-dependent modulation has become associated with proteins involved in oxidative stress regulation and maintaining metal-homeostasis within cells. To probe whether GRNs play a contributory role in such functions, we tested the metal binding potential of the reduced form of GRNs -3 and -5 under neutral and acidic pH mimicking cytosolic and lysosomal conditions, respectively. We found, at neutral pH, both GRNs selectively bind Cu(II) and no other divalent cations. Binding of Cu(II) also partly triggered the oxidative multimerization of GRNs via uncoordinated cystines at both pH conditions. Furthermore, binding did not induce gain in secondary structure and the protein remained disordered. Overall, the results indicate that GRN-3 and -5 have a surprisingly strong affinity for Cu(II) in the pM range, comparable to known copper sequestering proteins. This data also hints at a potential of GRNs to reduce Cu(II) to Cu(I), a process that has significance in mitigating Cu-induced ROS cytotoxicity in cells. Together, this report uncovers a metal-coordinating capability of GRNs for the first time, which could have profound significance in their structure and function.

## Introduction

Progranulin (PGRN), is an evolutionarily conserved protein observed widely in eukaryotes^1^. PGRN is a 68 kDa, glycosylated protein that is ubiquitously expressed in many cells and has pleiotropic roles in physiology such as neuronal growth, differentiation and survival^2-4^, wound healing and repair^5-6^, and immunomodulation^7^. The protein is also known to play a roles in tumor growth and metastasis by its growth-promoting and angiogenic characteristics^8^. PGRN is also linked to neurodegenerative disorders with the haploinsufficiency of the protein being implicated in frontotemporal dementia (FTD)^9^ while a complete loss of PGRN leads to neuronal ceroid lipofuscinosis, a lysosomal storage disease^10^. PGRN consists of seven and a half tandem-repeats of cysteine-rich modules called granulins (GRNs 1-7) and a partial para-GRN module. PGRN is secreted extracellularly and proteolytically cleaved by enzymes such as matrix metalloproteinase-12 (MMP-12)^11^, neutrophil elastase, and proteinase 3^12^ releasing the individual GRNs. In physiology, a variety of roles have been attributed to GRNs such as innate immune response^13^, neurotrophic roles^4^, while other studies ascribe pro-inflammatory functions, opposing that of the precursor^14^. Although the extracellular processing of PGRN and GRN formation is known^14-15^, recent studies have also revealed an intracellular localization of these modules. PGRN has been shown to be transported into the lysosome where it is processed by the cysteine peptidase cathepsin-L^16-17^ producing GRNs that are stable within the lysosomal environment^18^.

Individual GRNs (1-7) are 6 kDa proteins with a characteristic feature of unusually high cysteine content (∼20%)^1^, and have a consensus sequence of X_2-3_CX_5-_ 6CX_5_CCX_8_CCX_6_CCXDXXHCCPX_4_CX_5-6_CX, with the twelve conserved cysteines forming six putative disulfide bonds^19-20^. Studies into the structural properties of human GRNs have revealed that GRN-2, 4 and 5 adopt partially folded structure with a defined interdigitated disulfide bonding pattern, while the structures of other GRNs (1,3,6 and 7) are dominated by loops^20^. Previously, we have investigated the structure-function relationship of two GRNs; GRN-3^21-24^ and GRN-5^24^. Both GRNs show characteristics of classical intrinsically disordered proteins (IDPs) having no secondary structure especially in the fully reduced form^22, 24^. Abrogation of disulfide bonds in the reduced state renders the protein to be disordered and thermally unstable^22^. On the other hand, the disulfide-bonded oxidized GRN-3 showed high thermal stability but with a structure dominated by disordered loops^22^. Interestingly, both reduced and oxidized GRN-3 showed cellular activity implicating a possible redox-based regulation as a mode of functioning for GRNs^21, 23^.

Recently, the potential functional implications of disorder promoting sequences interspersed with order-promoting cysteines have begun to emerge. In eukaryotes, widespread presence of proteins, which are either conditionally disordered or contain intrinsically disordered regions are observed^25^. Such proteins display redox sensitivity by undergoing disorder-to-order under redox conditions and are implicated to play a role in plethora of biological processes. The ability within these cysteine-rich proteins to cycle between their redox states has recently been identified as a mechanism by which cells maintain homeostasis by buffering against potential redox stress^26-27^ The redox-sensitive regions are also often associated with metal binding; while proteins form disulfide bonds under oxidizing conditions (extracellular), metal binding is commonly observed under reducing conditions^25^. It is also noteworthy that the presence of redox-sensitive proteins increases with increasing organism complexity^25^. Furthermore, there exists a linear correlation between proteins containing disordered sequences and degree of cysteines in them in eukaryotes^28^. In the family of proteins containing more than 20% cysteines in the sequence, GRNs are somewhat similar to metallothioneins (MTs), which are metal sequestering proteins with crucial roles in maintaining metal-homeostasis in cells. MTs are ∼60 amino acid long proteins consisting of 20 cysteines (33%)^29^. MTs function as cellular reservoirs and scavengers of metal ions^29-32^ and have a prominent role in heavy metal detoxification^33-34^. Together, these observations combined with the fact that GRNs are known to be present extra and intracellularly with an abnormally high cysteine content, a redox-sensitive regulation of these proteins seems likely. Here we tested our hypothesis that GRNs bind divalent metal cations in reducing environments, in vitro. Results show that out of the seven metal ions tested, GRNs -3 and -5 specifically bind to Cu(II) in part via cysteine thiols and not the others. These results raise the question of whether copper binding is a key function of GRNs in their repertoire of biological function that has remained unnoticed thus far. The results presented here support this hypothesis and further elevates the relevance in their physiological functions.

## Materials and Methods

### Recombinant expression, purification, and generation of apoGRNs

GRN-3 and GRN-5 were recombinantly expressed and purified as previously described^22, 24^. Briefly, GRN-3 was expressed in SHuffle™ cells (New England Biolabs), while GRN-5 was expressed in Origami 2 DE3 (Invitrogen) as fusion constructs with a thioredoxin-A and hexa-histidine tag (TrxA-Hisx6-GRN). The constructs were purified using Ni-NTA affinity chromatography and eluate was treated with 5 mM ethylenediaminetetraacetic acid (EDTA) to chelate residual metal-ions. The buffers used in purification protocol were prepared using deionized water which was passed through a chelex resin column (Bio-rad) to exclude metal contaminants and obtain metal-free eluate. The glassware used in purification and buffer preparation were rinsed with chelex-treated water to wash residual contaminants. After exhaustive dialysis, fusion construct was cleaved with restriction grade thrombin (Bovine, BioPharm Laboratories) at 3U per 1 mg of the protein to remove both trxA and the His-tag. The reaction was incubated at room temperature (∼25°C) for 22-24 hours. Using a semi-preparative Jupiter® 5 µm 10×250 mm C18 reverse phase HPLC column (Phenomenex) protein was then purified by applying a gradient elution of 60 – 80 % acetonitrile containing 0.1% TFA on a UltiMate 3000 system (Thermo) as previously described^22^. The lyophilized apo-proteins were resuspended in either 20 mM HEPES-pH 7.0 or 20-mM ammonium formate-pH 4.5 prepared using chelex treated deionized water, as described before. Protein concentration was determined by measuring absorbance at 280 nm and using molar extinction coefficients of 6250 M^-1^cm^-1^ for GRN-3 and 7740 M^-1^cm^-1^ for GRN-5. Reduced forms of the proteins were used in experiments and were generated by incubating the proteins with a 20 stoichiometric excess of Tris(2-carboxyethyl)phosphine hydrochloride (TCEP) for 2 hours at RT.

### Metal-stock preparation for assays

Respective metal-stocks were prepared at a concentration of 2 mM, in either 20 mM HEPES-pH 7.0- or 20-mM ammonium formate-pH 4.5 in the presence of 12 mM glycine to avoid precipitation^35-36^. These buffers were prepared using deionized, chelex treated water, as described before and were stored at 4°C in dark.

### MALDI-ToF mass spectrometry

For determination of binding between metal-cations and GRNs, 1 mM metal-cations were incubated with the reduced, metal-free samples of 20 μM GRN-3 (MW 6367.7 Da) or GRN-5 (MW 6017.7 Da) in 20 mM HEPES buffer at pH 7.0 in the presence of 500 μM tris(2-carboxyethyl)phosphine hydrochloride Characterization of the protein-metal complexes was performed on a Bruker Datonics Microflex LT/SH ToF-MS system. For analysis of metal-protein reactions, 95.5 ng of GRN-3 and 90.2 ng of GRN-5 (15 pmoles) were spotted separately onto a Bruker MSP 96 MicroScout Target with a 1:1:1 ratio of sample:sinapinic acid matrix(saturated with acetonitrile and water): acetone. Instrument calibration was performed using Bruker Protein Calibration Standard I (Bruker Daltonics). Alkylation reactions were carried out by incubating respective metal-GRN samples prepared as described above, with 1mM iodoacetamide for 2h at room temperature. The samples were then prepared for analysis using MALDI-ToF-MS by spotting onto a Bruker MSP 96 MicroScout Target with a 1:1:1 ratio of sample:sinapinic acid matrix (saturated with acetonitrile and water): acetone.

### Fluorescence spectroscopy and determination of apparent dissociation constant

Intrinsic tryptophan fluorescence assays were performed on a Cary Eclipse spectrometer (Agilent Inc.) by exciting the samples at 285 nm and monitoring emission from 320 to 400 nm. 20 μM GRN (3 or 5) were titrated with increasing metal concentrations (10-240 μM) at pH 7.0 and pH 4.5 and spectra were recorded. Each spectrum represents an average of four repeat scan. The collected spectrum for each titration was normalized by integrating the area under the curve and the normalized intensities were plotted against concentration of Cu(II). The normalized curves were fitted to the one site binding equation:

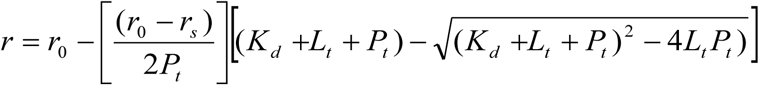

where *r*_*0*_ and *r*_*s*_ are fluorescence intensities in the absence and saturated levels of the ligand, Cu(II), respectively, while *L*_*t*_ and *P*_*t*_ are the respective total ligand (Cu(II)) and GRN concentrations and *K*_d_ represents the apparent dissociation constant 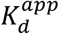. The data normalization and curve fitting were performed on Origin 8.5 graphing software. For obtaining the apparent dissociation constant with incorporation of dissociation constant of the, following equation was considered;

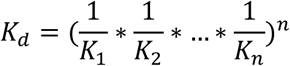

Using a value of 2.6 μM for the dissociation constant of glycine-Cu(II) complex^35-36^ and the values obtained for the respective GRN-Cu(II) complexes from intrinsic fluorescence assays, obtaining the final 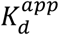 values in the picomolar range.

### Circular dichroism (CD) spectroscopy

Far UV-CD spectra were collected on Jasco J-815 instrument using a 0.1mm path length quartz cuvette (Precision cell). GRN samples incubated with respective buffers (20 mM HEPES pH 7.0 or 20 mM Ammonium formate) with or without Cu(II) as prepared previously for MALDI-MS characterization were suitably diluted from 20 μM to a concentration of 7 μM to avoid detector saturation at lower wavelengths. The samples were scanned from wavelengths 260 nm to 195 nm in the continuous scanning mode at a speed of 50nm/min. Each spectrum represents an average of three repeat scans.

## Results

### GRNs show redox sensitivity across their sequence similar to known metal binding proteins

The unique chemistry of cysteine residues marked by their high reactivity and an ability to form covalent bonds allows them to modulate the structure-function relation within proteins based on cellular cues. Such redox-responsive character has become associated with proteins that maintain cellular homeostasis, especially in response to oxidative stress or metallotoxicity^37^. To evaluate the potential redox sensitive character of GRNs -3 and -5, we utilized the IUPred2A computational platform^38^. IUPred2A computes and predicts the disorder propensities as well as redox sensitivity of a protein. As expected, analysis on MT-2, a well-known metal binding protein showed significant redox sensitivity across the entire stretch of the sequence (Fig 1 a). Analysis on GRNs revealed appreciable differences in the disorder propensities between the reduced and the oxidized forms (Fig 1b and c) for GRNs -3 and -5, respectively, and these are marked as regions of redox sensitivity (Fig 1; shaded purple areas) within the protein. IUPred2A analysis on β-defensin, a protein involved in anti-microbial functions used as a negative control, shows no redox sensitive regions in the protein despite some degree of disorder in the reduced form (Fig 1d). The disorder propensities of the free-thiol forms for MT-2 (Fig 1a), GRN-3 (Fig 1b) and GRN-5 (Fig 1c) have values between 0.75 and 1.0 indicating significant structural disorder. The redox sensitivity arises from both disorder within the sequence and due to a high number of cysteines interspersed within them^25^. In case of β-defensin, despite a ∼10% cysteine content, the two redox forms do not show significant differences in their disorder propensities leading to the prediction of a lack of redox sensitivity. This prediction correlates well with the known function of β-defensin as an anti-microbial peptide with no roles within a redox context^39^ and further underlines the importance of particular arrangement of cysteines within a sequence in imparting redox sensitivity. In MTs, two characteristic conserved motifs are utilized for metal coordination; xCCx and CxC (Fig 1a)^40^. The xCCx motif shares similarities with GRNs (Fig 1) and raises the potential for metal binding. These in-silico predictions align well with the previous findings on structural aspects of GRNs^21-22, 24^ and MTs^41-42^, where the reduced form of both proteins lacks structure.

**Figure 1:**
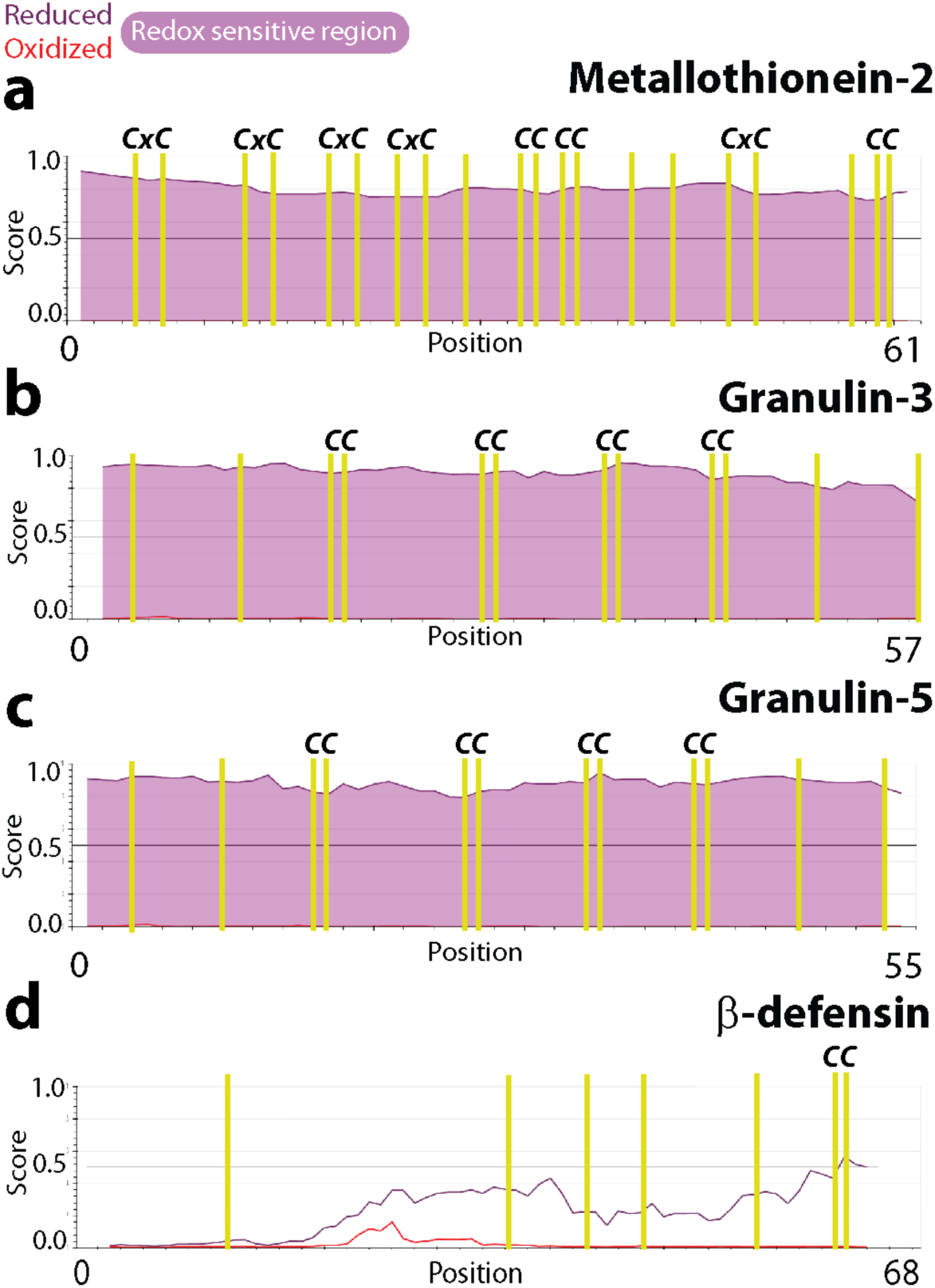
Redox sensitivity of cysteine-rich proteins predicted by IUPred2A. The disorder propensities of the redox forms of MT-2 (a), GRN-3 (b), GRN-5 (c) and β-defensin (d) are plotted with the disorder scores of the oxidized and reduced forms denoted by red and purple lines, respectively. Higher the score higher the disorder is. The positions of the cysteine residues are indicated by yellow vertical lines while the pink color between the red and purple lines denote the redox sensitive regions. MT-2 and β-defensins are shown as positive and negative controls, respectively.

### GRNs-3 and -5 bind Cu(II) but no other divalent metal cations

In order to see whether GRNs bind divalent metal cations, Cu(II) along with Ca(II), Co(II), Mg(II), Mn(II), Ni(II) and Zn(II) were used. These metals were chosen for their relevance in pathophysiology; Cu(II), Zn(II), Co(II), Mn(II) are cofactors for myriad enzymes, Ca(II) and Mg(II) are important in ionic balance and cellular signaling in addition to other important roles ^43^ and Ni(II) was chosen for testing the isoelectronic character and its involvement in toxicity^43^. These metal-ions were individually incubated in 50-fold molar excess with the reduced, metal-free samples of 20 μM GRN-3 (MW 6367.7 Da) or GRN-5 (MW 6017.7 Da) in 20 mM HEPES buffer at pH 7.0 in the presence of 500 μM tris(2-carboxyethyl)phosphine hydrochloride (see Methods). From these protein samples, aliquots of 15 pmols of GRN-3 and GRN-5 were then analyzed using MALDI-ToF MS (Fig 2; pH 7.0). At physiological pH of 7.0, both GRNs showed selective coordination up to 12 Cu(II) for GRN-3 and -5 (Fig 2a and b). GRN-3 showed a higher co-ordination than GRN-5; for GRN-3, seven and eight Cu(II) ions bound species were predominantly observed, while species with up to 12-Cu(II) ions bound was also evident. GRN-5 showed 1-Cu(II) bound species to be the most common one, with higher number of Cu(II)-ions bound species, with up to 11-Cu(II) bound forms, in an ascending order of abundance. GRN-3 showed more abundant Cu(II)-bound forms of the protein with only 10% remaining in the unbound, oxidized form (6357.8 Da) (Fig 2a), while GRN-5 bound Cu(II), the unbound, oxidized form (6000.7 Da) was ∼ 50% suggesting a diminished binding (Fig 2b). In contrast, no binding was observed for any of the other six divalent ions (Fig 2e, pH 7.0) with the exception of Zn(II); GRN-3 bound to one Zn(II) was ∼20% abundant while multiple Zn(II)-bound protein constituted ∼15% (Fig 2e). GRN-5 did not show the presence of Zn(II)-bound species. GRNs have recently been shown to be generated with the lysosomes also,^17-18^ while a central role of this organelle as a regulator of metal homeostasis has also come to the fore^44^. To mimic the binding of GRNs to metal-ions within a lysosomal environment, 20 μM GRN-3 or 5 was incubated with 50-fold molar excess of metal ions at pH 4.5, buffered with 20 mM ammonium formate. In the lysosome-like acidic environment, Cu(II) binding capability for GRN-3 and 5 were assessed by MALDI-ToF (Fig 2c and d). GRN-3 showed up to seven Cu(II) bound to the protein but with significantly decreased intensities as compared to those at pH 7.0 indicating diminished binding capability at the lower pH (Fig 2c). GRN-5 also showed significantly diminished binding with only one and three Cu(II) bound to the protein (Fig 2d). The relative abundance of the Cu(II)-bound protein was not more than ∼10% as compared to the apo form of the respective proteins, which showed ∼95% abundance based on the spectral intensities. A mass difference is observed for the non-coordinated species amongst the two pH conditions (pH 7.0 – pH 4.5) of both GRNs; 7.4 Da for GRN-3 (Fig 2a and c) and 4.1 Da for GRN-5 (Fig 2b and d) which can explained by the higher percentage of cysteines being present in the deprotonated thiolate form at the neutral pH^45^. Both proteins also showed inability to binding other metal cations at pH 4.5 (Fig 2e; pH 4.5). The lack of metal binding at low pH is not surprising and could likely be due to chelating functional groups such as thiols being predominantly present in a protonated form^45^.

**Figure 2:**
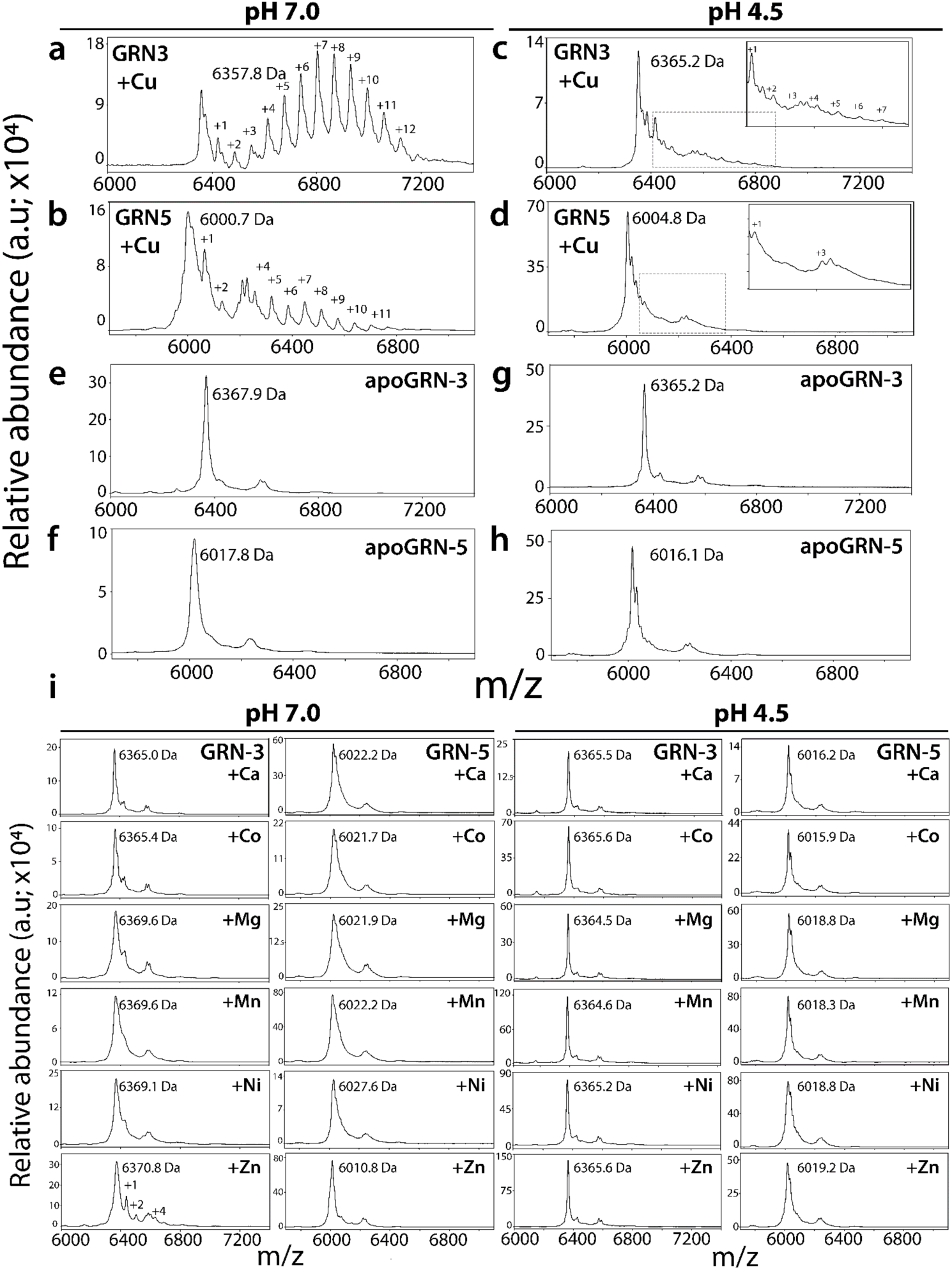
Metal binding analysis of GRNs by MALDI-ToF. GRN-3 or GRN-5 (20 μM) were incubated separately with a 1mM (50-fold molar excess) of respective metal cations in 20 mM HEPES, 12 mM glycine at pH 7.0 or in 20 mM ammonium formate, 12 mM glycine at pH 4.5 individually, at 4°C overnight. For MALDI-ToF-MS analysis, aliquots containing 6 x10^3^ pmol/L of GRN-3 and GRN-5 from the respective reactions loaded on to the plate. Cu(II) binding to GRN-3 (a) or GRN-5 (b) at neutral pH, and to GRN-3 (c) or GRN-5 (d) at pH 4.5. Apo GRN-3(e, g) or apo GRN-5 (f, h), prepared in similar buffers at the respective pH but without the metal cations. i) Spectra of Ca, Co, Mg, Mn, Ni and Zn binding to GRN-3 or GRN-5 at pH 7.0 (left panel) and at 4.5 (right panes).

Based on our hypothesis, free thiols in cystines are involved in metal coordination. Therefore, to determine how many cystines are involved in coordination, alkylation of free thiols in the Cu(II)-bound holoproteins was conducted with iodoacetamide (IAA) for both GRNs. IAA reacts with free thiols to forms acetamide adduct (58 Da) with sulfur^46^. The Cu(II)-bound GRN-3 at pH 7.0 displayed a gaussian distribution centered around ∼two alkylated species (6490.8 Da) suggesting that most of the cystines were coordinated to Cu(II) at pH 7.0 (Fig 3a). Heterogeneous population of species involving metal-bound and alkylated, only alkylated, or only metal-bound forms, should appear as clustered signals at m/z values of ∼6700-7000 Da. Clearly, the gaussian curve is unsymmetrical towards the trailing edge suggesting this is indeed the case (Fig 3a). GRN-5 at the same pH showed no free thiols available for alkylation by IAA with a major peak of 6009.0 Da corresponding to the non-alkylated, oxidized form (Fig 3b). Similar to GRN-3, a well-defined non-alkylated signal is followed by trailing peaks indicating the presence of multiple alkylated, metal bound forms with a descending level of abundance (Fig 3b). Furthermore, despite attenuated binding of Cu(II), both GRNs under acidic conditions also show decreased alkylation suggesting that the interaction with the metal-ion leads to the unavailability of free thiols within GRN-3 (Fig 3c) and GRN-5 (Fig 3d), possibly due to induced oxidation (further elaborated in Discussion). The spectra of GRN-3 showed a defined signal at 6464.4 Da which corresponds to the monomeric protein without any alkylation (Fig 3c). Similar spectra is observed for GRN-5, with a signal at 6014.9 Da (Fig 3d), again indicating the lack of thiols available for alkylation. The observation also suggests potential oxidation of cysteines induced by Cu(II) at low pH. The alkylation of GRN-3 incubated with other metal ions at pH 7.0 indicated most of the thiols were free and available for labeling by IAA, except for the sample with Mn(II) which showed a decrease of two free thiols, while the protein with Zn(II) displayed a heterogenous mixture of multiply-alkylated forms; with four, seven and eleven free thiol species (6569.9 Da, 6738.1 Da, 7013.7 Da; Fig 3e, GRN-3 at pH 7.0) being the predominant ones.. In the presence of Zn(II) ion, a similar spectrum containing multiple alkylated species is observed for GRN-5, with four and eleven free thiols forms (6240.9 Da, 6660.1 Da; Fig 3e, GRN-5 at pH 7.0). These observations along with the detectable binding of GRN-3 to Zn(II) (Fig 2e, GRN-3 at pH 7.0) further indicates potentially weak coordination of Zn(II) by GRNs at neutral pH. In acidic pH, GRNs show decreased availability of thiols for alkylation as indicated by the spectra of GRN-3 and 5 control samples without any metal-ions (Fig 3e, GRN-3 and GRN-5 at pH 4.5). The lower pH increases the concentration of the protonated thiols and hence, decreases alkylation by IAA, an observation that is in line with a decrease in rate of thiol alkylation by 2-vinylpyridine in an acidic pH^47^ Compared to the control reactions, no other metal-ion incubated samples showed any significant deviation in terms of alkylation patterns, except for Zn incubated samples, where both proteins showed an increase of two-three thiols available for alkylation (Fig 3e, GRN-3 and GRN-5 at pH 4.5). Alkylation of thiols using IAA is widely performed, especially due to low levels of unwanted, side reactions that accompany this chemistry^46,^ 48. Despite this, our results indicate the presence of alkylation of residues other than cysteine, as observed in Fig 3e, GRN-3 at pH 7.0, where a discernible signal from a species with 13-alkylated functional groups is present at a mass of 7121.3 Da. This points towards alkylation of either an acidic residue or histidine, along with cysteine thiols^46^. Another side reaction observed is the modification of N-terminal methionine residues of GRNs on reaction with the iodine containing alkylating agent resulting in the loss of a 47 Da fragment from these proteins^49^. This loss is evident in the IAA reactions of GRN-5 (Fig 3e, GRN-5 at pH 7.0 and pH 4.5) where a signal is observed at 5970.7 Da, preceding the monomeric, non-alkylated form of the protein (GRN-5 MW; 6017.7 Da).

**Figure 3:**
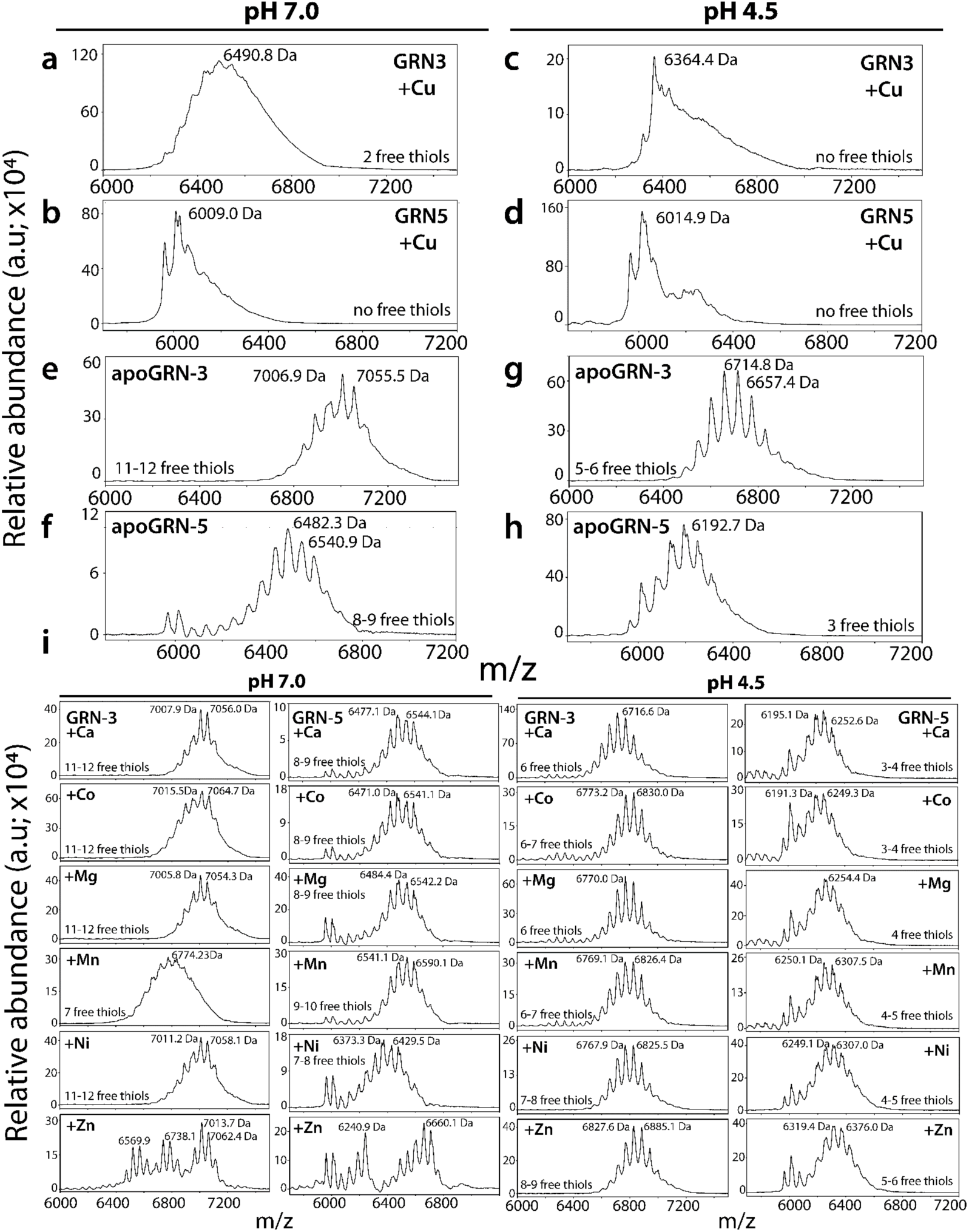
Determination of free thiols by IAA alkylation of GRNs. Holoproteins from Fig 2 were used for alkylation reactions using 1 mM iodoacetamide (IAA) and aliquots of samples were analyzed using MALDI-ToF-MS. The spectra of alkylated samples of Cu(II) incubated GRN-3 (a) and GRN-5 (b) in a pH 7.0 buffer and GRN-3 (c) and GRN-5 (d) in pH 4.5. e-f) MALDI-ToF-MS spectra of apoGRNs at the two pH conditions tested. i) The spectra of alkylation reactions of GRN-3 and -5 incubated with other metal-ions (Ca, Co, Mg, Mn, Ni and Zn) at neutral and acidic pH.

### Copper binding induces multimerization of GRNs

As mentioned earlier, the interaction of GRNs with Cu(II) decreased the availability of free thiols for alkylation suggesting their potential oxidation (Fig 3a-d). To determine whether the thiols are involved in intra or inter-molecular disulfide bonding following their interaction with Cu(II) and other metal-ions, we subjected the samples from Fig 2 to polyacrylamide gel electrophoresis (PAGE). At neutral pH, GRN-3 (MW; 6367.7 kDa) is observed to undergo multimerization in the presence of Cu(II) (Fig 4a, +Cu). Band corresponding to the dimeric form (MW; 12.6 kDa) is prominently visible while faint bands corresponding to monomeric and trimeric species (MW; 19.1 kDa) are also observed (Fig 4a). On the other hand, only a band corresponding to the monomeric form is seen in the presence of other metal cations, similar to apoGRN-3 (Fig 4a). Prior characterization of GRN-5 (MW; 6017.7 Da) in our lab revealed the monomeric protein displays a decreased electrophoretic mobility in PAGE, a commonly observed property among disordered proteins^24, 50^. In line with our previous observation, apoGRN-5 is observed as a band at ∼30 kDa at neutral pH (Fig 4b, apoGRN-5). In the presence of Cu(II), a diffuse, ‘smear’ pattern is visible with no distinct band, indicating significant multimerization of the protein (Fig 4b, +Cu). All other metal cations, with the exception of Zn(II) and Ni(II), showed only a single band similar to aopGRN-5 (Fig 4b). In the presence of Zn(II) and Ni(II), bands are observed at ∼17 and 20 kDa, respectively (Fig 4b). It is unclear as to why such an odd migration is observed for these metals. Nevertheless, multimerization is not observed with other metal cations in a manner that is observed with Cu(II). At pH 4.5, all metal-incubated GRN-3 samples show the presence of a distinct trimer at 18 kDa (Fig4c). In the presence of Cu(II), the degree of multimerization is prominent with a heterogenous mixture of multimeric species forming a ‘smear’ pattern, where the monomeric, dimeric and trimeric bands are still discernible (Fig4c, +Cu). Other metal-ions failed to induce the multimerization of GRN-3, similar to the observation at pH 7.0 (Fig 4c). At pH 4.5, GRN-5 shows an electrophoretic pattern similar to one observed at neutral pH (Fig 4b and d). The presence of Cu(II) induces multimerization of GRN-5, it is not as pronounced as that observed at pH 7.0 (Fig 4b and d, +Cu). A distinct band is observed at ∼30 kDa, similar to apoGRN-5 and a characteristic smear of multimerized protein extending above 50 kDa. Sample with Zn(II) shows higher electrophoretic mobility in the acidic regime as well, with the protein band present at ∼17 kDa. At both the pH conditions tested, Mg(II) and Mn(II) are observed to induce a visible smearing of GRN-5 towards the lower molecular weight region, indicating higher electrophoretic mobility of the species formed (Fig 4b and d, +Mg, +Mn). These results indicate that interaction with Cu(II), but not necessarily binding, seems to induce the oxidation of cysteines within GRNs with consequent multimerization of the protein.

**Figure 4:**
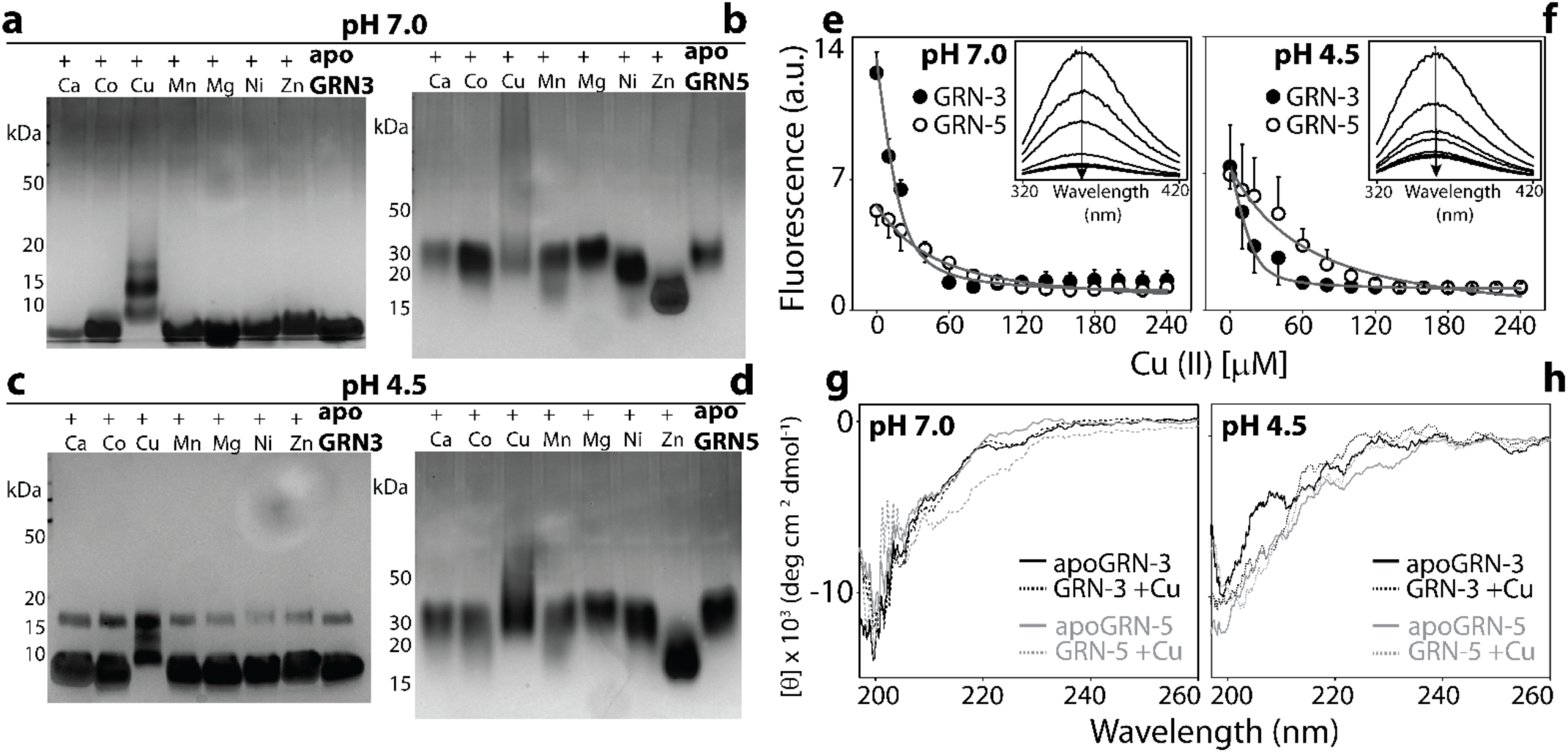
Binding and metal-induced structural changes in GRNs. Metal-incubated samples of GRNs were subjected to PAGE in non-reducing conditions. (a-d) and spectroscopic analysis (e-h). Silver stained gels of Apo and holoGRN-3 (a) and apo and holoGRN-5 (b) at neutral pH; apo and holo GRN-3 (c) and GRN-5 (d) at pH 4.5. Intrinsic tryptophan fluorescence of GRNs (20 μM) measured as a function of increasing Cu(II) (10-240 μM) concentration at pH 7.0 (e) and pH 4.5 (f) to evaluate tertiary structural changes. (inset): raw spectral scans shown as a function of increasing Cu(II) concentrations (arrow). g and h) Far-UV circular dichroism (CD) spectra of apo and holoGRNs incubated with Cu(II at neutral pH (g) and at pH 4.5 (h).

To see whether metal binding induces changes to the conformational environment surrounding the lone tryptophan residue in the proteins, the reduced apo forms of 20 μM GRN-3 or GRN-5 were titrated with increasing molar concentrations (10-240 μM) of Cu(II) and the intrinsic tryptophan fluorescence was measured. This informs about the changes in the environment around the tryptophan residue as a function of metal ion concentration, and therefore allows the calculation of apparent binding affinity for the metals^21, 36, 51-52^. At neutral pH, GRN-3 shows a higher intrinsic fluorescence than GRN-5 in absence of Cu(II) (Fig 4e). Such a variation in fluorescence intensities suggests an inherent difference in the tryptophan environment amongst the two proteins^52^. Increasing Cu(II) concentrations leads to the decrease in tryptophan fluorescence intensity in both GRNs (Fig 4e). This observation indicates an increased solvent exposure upon metal binding ^22^. The data was fitted to a one-site binding equation (detailed in Methods) to obtain an apparent dissociation constant 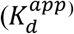 for GRNs and Cu; 4.0 ± 1.5 μM for GRN-3 and 35.4 ± 8.2 μM for GRN-5. This 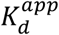 however, does not account for the binding of Cu(II) by glycine used in the buffer^35-36^. Upon correcting the binding constant for this non-specific competitive binding of glycine (see Methods), 10.4 pM and 92.0 pM for GRN-3 and GRN-5, respectively are obtained. At acidic pH, both GRNs have equivalent fluorescence intensities in absence of Cu and show a similar trend of attenuation of fluorescence intensity with increasing Cu(II) concentration. The apparent corrected dissociation constants for this interaction are 6.9 pM for GRN-3 and 130.5 pM for GRN-5. These apparent dissociation constants are comparable to those of known Cu-coordinating proteins, such as MT-2^53^. It has to be borne in mind these values only reflect indirect affinities for Cu; at pH 4.5, no actual binding is observed between Cu and GRNs (Fig 2c and d), and yet quenching of fluorescence is pronounced. This anomaly indicates that the dissociation constants inform us about the structural changes around tryptophan, which may arise not only due to direct metal binding but also due to consequential multimerization.

Intrinsically disordered proteins have been known to obtain structural order upon metal coordination or binding to ligands^50^. To probe whether metal binding induces secondary structural changes within GRNs, far-UV circular dichroism (CD) was used samples from Fig 2, which were suitably diluted to 7 μM GRN concentrations. In agreement with previous biophysical studies on GRN-3 and GRN-5, at pH 7.0, both apo proteins display spectra characteristic of random coil with a minima observed at 200 nm (Fig 4g). Cu-binding fails to induce any secondary structural elements within GRN-3 as evidenced by a spectrum similar to its apo form. At neutral pH, GRN-5 in presence of Cu shows a weak minimum at ∼215 nm, indicative of some β-sheet content within the protein. Such induction of secondary structure within unstructured proteins by metal coordination is well known, although the intensity of the ellipticity observed here suggests GRN-5 remains mostly unstructured. At low pH also, apo forms of GRN-3 and GRN-5 display a random coil spectra, with a minimum at 199nm, indicating these proteins remain unstructured in the acidic environment (Fig 4h). As would be expected, the Cu(II)-incubated samples show overlapping signals with the apo-proteins (Fig 4h), since no coordination is detected with Cu(II) at this pH (Fig 2 c and d). These results show that interaction with Cu(II) brings about tertiary and quaternary structural changes within GRNs, as evidenced by the fluorescence quenching (Fig 4e and f) and multimerization (Fig 4a-d), respectively, but fails to induce significant secondary structural changes. These observations are similar to those observed for MTs, where these proteins do not attain any secondary structure on metal coordination but become conformationally constrained^40, 42, 54^.

## Discussion

The metal-coordinating potential of GRNs has been hypothesized since their initial characterization^55-56^ while the recent study by Fang and colleagues strengthened the rationale for such an investigation by indicating potential coordination of metals by GRNs^57^. Our report brings forth a functionality of GRNs that has remained largely unexplored, particularly at the molecular level. Properties such as the unusually high number of conserved cysteines and disorder within the sequence in the fully reducing conditions^21^ make GRNs similar to MTs, which are known to bind Cu(II), Zn(II) and Cd(II) ions, among others^40^. Further motivated by the IUPred2A predictions that show GRNs to be similar to MTs in redox sensitivity, we set out to test whether GRNs show affinity for divalent cations and reveal potential promiscuity or specificity of such metal coordination^57^. The results indicate that both GRNs -3 and -5 have affinity towards Cu(II) with GRN-3 showing greater affinity than GRN-5 at neural pH. However, both GRNs, at acidic and neutral pH, failed to bind any of the other divalent metal cations, highlighting a selectivity towards Cu(II).

We also observe that binding of copper fails to induce any secondary structural changes in either GRN-3 or GRN-5, a characteristic similar to that of MTs^54^. Such conformational flexibility has become an identifiable property of IDPs that are often associated with pleiotropic roles^50, 58-59^. These observations align well with the multiple functional roles that GRNs are known to participate in and suggest copper-binding could be yet another function for these protein modules. Furthermore, interaction with copper, and not other metals, induces the formation of multimeric GRN-3 or -5, under both pH conditions tested. Alkylation experiments using IAA suggest that the interaction with copper leaves no or few free cysteines within both GRNs. A majority of cysteines are either coordinated to copper or involved in intra- or inter-molecular disulfide bond formation, or both. This brings to light two possible scenarios that may contribute to the mechanism of GRN functions: *i*) A fraction of Cu(II) oxidizes thiols in some cysteines to form disulfide bonds leading to multimerization and the protein may bind either Cu(I) (generated from the reduction of Cu(II)) or Cu(II) (present in excess), or *ii*) Cu(II) coordinates to cysteines from two GRN monomers forming a bridge ligand leading to multimerization. The former scenario is supported by the fact that decreasing the pH to 4.5 decreases the extent of multimerization of GRNs due to the protonation of thiols. The results here show the coordination of Cu(II) at neutral pH, and its abrogation in an acidic pH, provides cues into their potential cellular roles; GRNs could mitigate cu-induced cytotoxicity by binding Cu(II) in the reducing, cytoplasmic environment and releasing them within the acidic, lysosomal milieu where GRNs are known to localize ^17-18^. Secondly, the dissociation constants of Cu(II) with GRNs measured indirectly by intrinsic tryptophan fluorescence as a function of metal concentrations reveal that the affinities both GRNs for Cu(II) are in the picomolar range. This is comparable to the affinity of MTs for Cu(II), which is in the pM-fM range^60^. Mammalian cells mainly employ MTs and superoxide dismutases (SOD) to protect themselves from ROS activation by excess Cu(II) ions. The comparable affinity range of copper for GRNs, MTs and SODs as well as the potential reduction of Cu(II) to Cu(I) by the cysteines, suggest that GRNs may play a protective role during Cu(II) induced oxidative stress to the cells similar to MTs and SODs by either promoting disulfide bonds or binding to free cystines or both. However, unlike MTs and SODs, GRNs seem to exhibit selectivity towards copper based on the results obtained here. What precise physiological significance copper binding holds is unclear at this point but one can speculate that a network of copper binding proteins might be employed, driven by affinity gradients^53^. Intracellular GRNs may participate in mitigating ROS toxicity by behaving as an additional metal scavenger which may be needed during acute Cu(II) influx. Such a mechanism can bypass the ROS generation induced by free Cu(II) and its associated Fenton-like chemistry. To our knowledge, this study is the first to uncover the Cu-binding ability of GRNs, adding it to the known functional repertoire of these proteins. Although intriguing, these results warrant further investigations into the interactions of GRNs with metal-cations to obtain a clear picture of what roles GRNs play in metal homeostasis in norm and pathology.

## Acknowledgements

The authors would like to thank the following agencies for financial support: National Institute of Aging (1R56AG062292-01) and the National Science Foundation (NSF CBET 1802793) to VR. The authors also thank the National Center for Research Resources (5P20RR01647-11) and the National Institute of General Medical Sciences (8 P20 GM103476-11) from the National Institutes of Health for funding through INBRE for the use of their core facilities.

## Notes

### Competing Interest Statement

The authors have declared no competing interest.

